# Suppressing the morning cortisol rise after memory reactivation at 4 a.m. enhances episodic memory reconsolidation in humans

**DOI:** 10.1101/2020.11.30.404707

**Authors:** Despina Antypa, Aurore A. Perrault, Patrik Vuilleumier, Sophie Schwartz, Ulrike Rimmele

**Affiliations:** Laboratory of Neurology & Imaging of Cognition, Department of Basic Neurosciences, University of Geneva, Switzerland; Swiss Center of Affective Sciences, University of Geneva, Switzerland; Department of Psychiatry and Behavioral Sciences, University of Crete, Greece; Laboratory of Sleep and Cognition, Department of Basic Neurosciences, University of Geneva, Switzerland; Sleep, Cognition and Neuroimaging Laboratory, Department of Health, Kinesiology and Applied Physiology, Concordia University, Montreal, QC, Canada; Faculty of Psychology and Educational Sciences, University of Geneva, Switzerland; Center for Interdisciplinary Study of Gerontology and Vulnerability (CIGEV), University of Geneva, Switzerland

## Abstract

Evidence from animal and human research shows that established memories can undergo changes after reactivation through a process called reconsolidation. Alterations of the level of the stress hormone cortisol may be one way of manipulating reconsolidation. Here, in a double-blind, within-subject design, we reactivated a 3-day-old memory at 3:55 a.m., immediately followed by oral administration of metyrapone vs. placebo, to examine whether metyrapone-induced suppression of the morning cortisol rise may influence reconsolidation processes during and after early morning sleep. Crucially, reactivation followed by cortisol suppression vs. placebo resulted in enhanced memory for the reactivated episode (tested four days after reactivation). This enhancement after cortisol suppression was specific for the reactivated episode vs. a non-reactivated episode. These findings suggest that when reactivation of memories is immediately followed by suppression of cortisol levels during early morning sleep, reconsolidation processes change in a way that leads to the strengthening of episodic memory traces.

## Introduction

You wake up in the middle of the night. Just a hint of a past memory comes through your mind and this may be enough to change this memory. Several studies have reported that the mere reactivation of a memory can change the way this memory is stored in the brain, through a process described as reconsolidation (*1–6*). Reconsolidation has been proposed to be an additional memory stage, when formed memories become prone to change, after the reactivation of their established memory trace (*7, 8*). In particular, targeted pharmacological and behavioural manipulations following memory reactivation are thought to modulate the reconsolidation process and thus critically change an already formed memory (*9–14*).

The stress hormone cortisol in humans (corticosterone in rodents) has been proposed to modulate the reconsolidation process (*15, 16*). If a stressor follows memory reactivation, the reactivated memory is changed at a later test, advocating for a modulation of reconsolidation by stress-induced increases in cortisol levels (*17–20*). Pharmacological studies support that cortisol plays a critical role in memory reconsolidation (*15, 16*). Fear memories can be altered in long-term in animals and humans, when after their reactivation, corticosterone or cortisol is pharmacologically administered (*21–25*). This cortisol-induced alteration of reconsolidation is possibly dose-dependent: For example, when stress-induced corticosterone levels are lowered with metyrapone after fear memory reactivation, then the typically stress-induced fear memory disruption during reconsolidation is reversed (*26*).

Interestingly, cortisol levels do not only change pharmacologically or upon encountering a stressor, but also vary physiologically throughout a 24-h day/night cycle: Following a circadian rhythm, cortisol levels decrease in the evening and during early sleep, and rise again in the early morning, leading to a robust morning cortisol peak at the time of waking up (*27*). Previous studies have shown that memory consolidation during sleep depends on this physiological early-night inhibition of cortisol release co-occurring with a distinct sleep-pattern (*28–30*). In particular, the decrease in cortisol levels as it naturally occurs in the first half of the night accompanied by long blocks of slow-wave sleep (SWS), has been proposed to enhance consolidation of hippocampus-dependent memories (such as memory of episodes). In contrast, the physiological morning cortisol rise in humans, starting around 4 a.m. in the morning, accompanied by key changes in sleep patterns (shorter blocks of SWS and longer blocks of REM sleep; *31*) has been suggested to hinder the consolidation of newly encoded memories, possibly by interrupting the transfer of information between hippocampus and prefrontal cortex (*32, 33*). Analogously to consolidation, reconsolidation processes have been reported to be susceptible not only to cortisol (e.g., as described above, by: *13, 24*) but also to sleep manipulations (*34*). Thus, not only consolidation but also reconsolidation processes may be affected by the interaction of the physiological morning cortisol rise and its associated sleep patterns.

To test this possibility, here we examined episodic memory reconsolidation taking place during the physiological morning cortisol rise vs. when the morning cortisol rise was pharmacologically suppressed using a within-subject crossover design (N = 20). We administered metyrapone at 4 a.m. in the morning, as we did in a previous study that showed robust suppression of the morning cortisol rise (*35, 36*). We combined this morning cortisol suppression with a previously established reconsolidation paradigm (*11, 13, 14*) to test whether memory reactivation at 3:55 a.m. immediately followed by cortisol suppression changes reconsolidation, hence resulting in altered later memory of the reactivated episode. For the reactivation of a consolidated episodic memory, we used a reminder cue that was followed either by physiological cortisol rise (placebo condition) or pharmacologically suppressed cortisol levels (metyrapone condition) during and after early morning sleep. In particular, two days after the encoding of two stories, we re-activated the consolidated memory of one of the two stories with a reminder cue at 3:55 a.m. and thereafter administered either placebo or the cortisol synthesis inhibitor metyrapone in a double-blind design. Pill administration was followed by sleep until 6:45, for which we assessed standard polysomnographic (PSG) recordings. Four days later, we tested memory for both the reactivated and the non-reactivated story. We expected that reactivation followed by a normal physiological morning cortisol rise would disrupt reconsolidation, in analogy to impairing effects of stress induction on reconsolidation and of morning cortisol rise on consolidation (*20–23, 37, 38*). Moreover, we put forward the hypothesis that cortisol suppression can reverse this effect, by enhancing reconsolidation processing.

## Results

### Post-reactivation cortisol suppression enhances episodic memory reconsolidation

Cortisol suppression at 4:00 a.m., directly after memory reactivation, enhanced memory performance in a multiple-choice recognition memory task assessed four days after re-activation [main effect of substance: *F*(1,17) = 6.395, *p* =.022, η^2^ = .273; *M*_*Metyrapone*_ = .51, *SE* = .03 vs. *MPL =* .45, *SE* = .02; **Figure 1A**].

**Figure 1.**
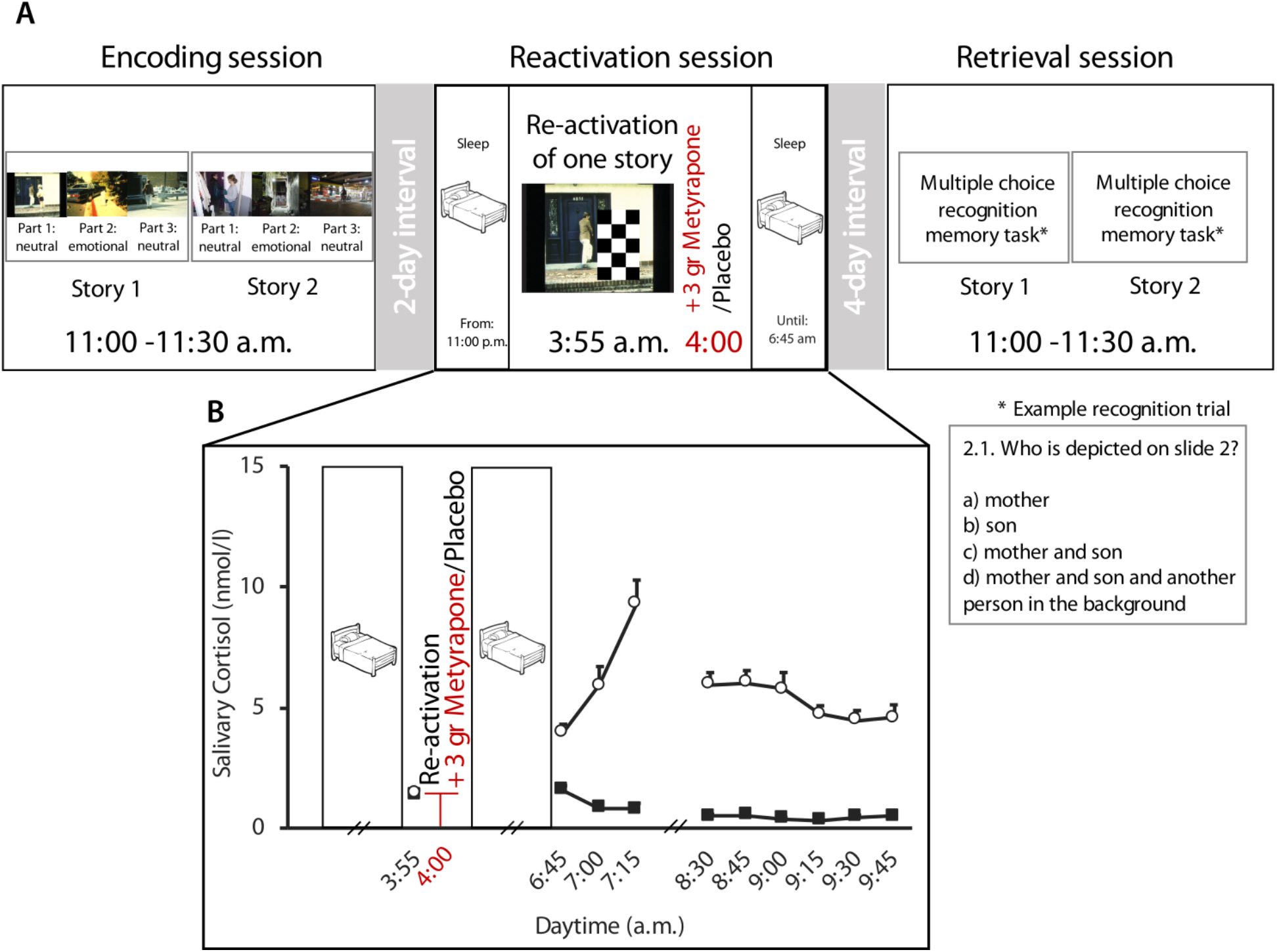
Experimental procedure (A) and cortisol levels (B). (A) Each participant was tested once in a metyrapone and once in a placebo condition that both comprised three sessions. The order of conditions was counterbalanced across participants. Both conditions comprised three sessions (Encoding, Reactivation, Retrieval). At the Encoding Session, participants were presented two stories. At the Reactivation Session (two days after encoding), participants slept in the laboratory. After awakening at 3:55 a.m., one of the two stories was reactivated and metyrapone or placebo was administered at 4:00 a.m. They then slept again until 6:45 a.m. At the Retrieval Session (seven days after encoding), memory was tested for both the reactivated and the non-reactivated story with a multiple-choice recognition memory task. (B) Cortisol levels: Mean ± SE salivary cortisol concentration for the Reactivation Session. Baseline cortisol levels prior to reactivation and substance administration (at 3:55 AM) did not differ between conditions. Metyrapone (black squares) suppressed cortisol levels for all other measurement points.

Most importantly, there was a substance by reactivation interaction [*F*(1,17) = 4.678, *p* =.045, η^2^ = .216]: memory performance for the reactivated story was significantly higher in the metyrapone condition (*M*_*Metyrapone*_ = .55, *SE* = .04) in comparison to the reactivated story in the placebo condition [*M*_*PL*_ = .45, *SE* = .02; *t*(17) = 3.817, *p* =.001, *d* = .890]. Crucially, in the metyrapone condition, memory was higher for the reactivated (*M*_*RS*_ = .55, *SE* = .04) in comparison to the non-reactivated story [*M*_*NRS*_ = .47, *SE* = .03; *t*(17) = 2.578, *p* =.020, *d* = .608]. There was no difference in memory performance for the non-reactivated stories between the metyrapone vs. placebo conditions [*t*(17) = .488, *p* = .632], and no difference between reactivated and non-reactivated stories in the placebo condition [*t*(17) = −.097, *p* = .924; **Figure 2A**]. Lastly, there was no main effect of reactivation [*F*(1,17) = 3.019, *p* =.100].

**Figure 2.**
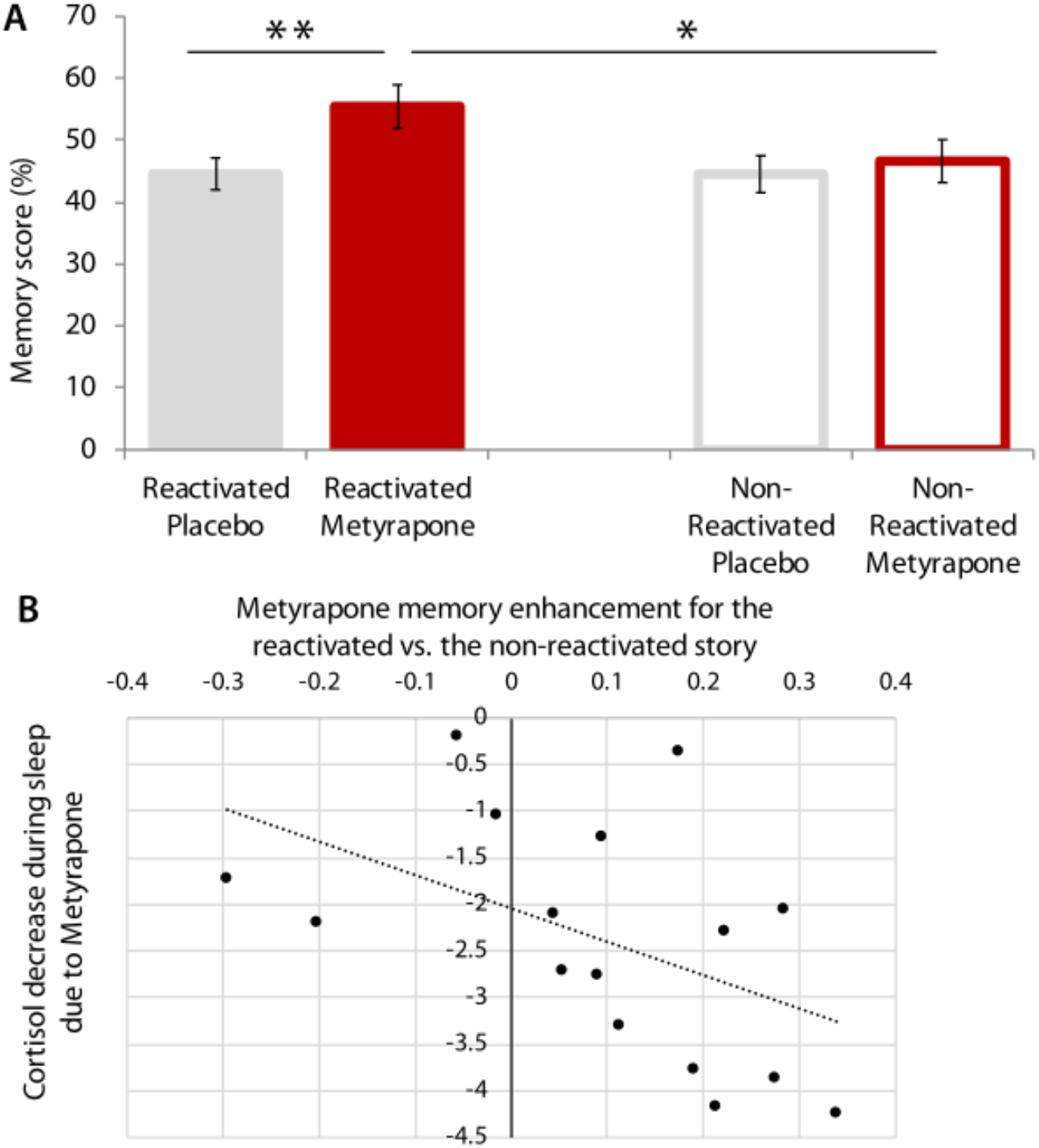
Memory performance in the metyrapone vs. placebo condition of reactivated vs. non-reactivated story (A) and relation between individual differences in memory performance and cortisol suppression during sleep (B). (A) Pharmacologically suppressing cortisol at 4:00 a.m. directly after re-activation of a story at 3:55 a.m. enhanced memory performance for the re-activated story four days later in the metyrapone vs. placebo condition. Importantly, cortisol suppression resulted in enhanced memory only for the reactivated but not the non-reactivated story (significant substance by reactivation interaction). Error bars indicate SE and * *p* < .05, ** *p* < .01. (B) Individual metyrapone memory enhancement for the reactivated vs. non-reactivated story was negatively correlated with the individual cortisol decrease due to metyrapone during sleep (*τ* = −.450, *p* = .015).

Individual metyrapone memory enhancement for the reactivated vs. non-reactivated story was negatively correlated with the individual cortisol decrease due to metyrapone <u>during</u> sleep (*τ* = −.450, *p* = .015; **Figure 2B**). In contrast, there was no correlation between metyrapone memory enhancement for the reactivated story and cortisol decrease due to metyrapone <u>after</u> sleep (*p* = .805).

### Metyrapone administration suppresses morning cortisol rise

Prior to reactivation and prior to substance administration (at 3:55 a.m.), baseline cortisol levels were comparable between the placebo (placebo baseline: *b* = .071, *t*(393) = 1.067, *p* = .287) and the metyrapone condition [metyrapone baseline: *b* = .614, *t*(393) = −.504, *p* = .614]. Following reactivation and substance administration, cortisol levels were lower after metyrapone vs. placebo administration [main effect of substance: *F*(1,373) = 1321, *p* <.001; substance by time interaction: *F*(10,373) = 19.584, *p* <.001; main effect of time: *F*(10,374) = 6.988, *p* <.001] for all measurements taken between 6:45 and 10:00 a.m. (all *p* < .001). The maximum difference to baseline was observed at 7:15 [placebo: *b* = .861, *t*(393) = 10.263, *p* < .001; metyrapone: *b* = −1.161, *t*(393) = −9.679, *p* < .001; see Table S1; **Figure 1B**].

### Metyrapone administration alters subsequent sleep period

As expected, before substance administration, the metyrapone and the placebo condition did not differ in sleep duration (measured by total sleep period, TSP, and total sleep time), or in the proportion of time spent in the different sleep stages during the first part of the night (i.e. from sleep onset to 3:55 a.m.) between (all *p* > 0.1; see **Table 1**). However, metyrapone intake at 4:00 a.m. significantly affected the subsequent sleep period. Compared to placebo, metyrapone increased the time spent awake between 4:05 a.m. and 6:45 a.m. by approximately 15 minutes (from 5% to 18% of TSP) in comparison to the placebo condition [*t*(10) = 3.952, *p* = .003, *d* = 1.192]. In addition, metyrapone altered the proportion of time spent in different sleep stages as revealed by an increase in N1 duration [*t*(10) = 4.953, *p* = .001, *d* = 1.493], and a decrease in N3 [*t*(10) = 4.238, *p* = .002, *d* = 1.278] and REM duration [*t*(10) = 4.630, *p* = .001, *d* = 1.396; see **Table 1).** Note that metyrapone intake did not affect the duration of N2 ([*t*(10) = .1704, *p* = .868]. The increased time spent awake after substance administration also affected total sleep time, which was reduced by 11%, and consequently decreased sleep efficiency (*M*_*PL*_ = 94.24 ± 5.1, *MM* = 81.87 ± 7.5; [*t*(10) = 3.952, *p* = .003, *d* = 1.192] during the second part of the night (i.e., after substance administration: from 4:05 a.m. to 6:45 a.m.).

**Table 1.** Sleep architecture

|  | Before substance administration<br>Sleep onset to 3:55 a.m. |  |  |  | After substance administration<br>4:05 a.m. to 6:45 a.m. |  |  |  |
| --- | --- | --- | --- | --- | --- | --- | --- | --- |
|  | Placebo | Metyrapone | *t*(10) | *p* | Placebo | Metyrapone | *t*(10) | *p* |
| TSP<br>(min) | 271.63<br>± 19 | 269.45<br>± 31 | .216 | 0.834 | 143.31<br>± 15 | 147.18<br>± 4 | .758 | 0.466 |
| TST<br>(min) | 262.5<br>± 20 | 258.18<br>± 24 | .486 | 0.638 | 135.36<br>± 19 | 120.54<br>± 12 | 1.976 | 0.076 |
| N1<br>%TSP | 2.24<br>± 1.29 | 3.16<br>± 2.01 | 1.370 | 0.201 | **7.71**<br>± **6.71** | **29.52**<br>± **18.77** | **4.593** | **0.001** |
| N2<br>%TSP | 37.38<br>± 9.44 | 39.23<br>± 6.82 | .732 | 0.481 | 41.14<br>± 11.98 | 40.08<br>± 18.71 | .170 | 0.868 |
| N3<br>%TSP | 38.87<br>± 8.16 | 38.48<br>± 6.73 | .128 | 0.900 | **12.95**<br>± **10.21** | **0.13**<br>± **0.43** | **4.238** | **0.002** |
| REM<br>%TSP | 18.11<br>± 5.61 | 15.16<br>± 4.05 | 1.471 | 0.172 | **32.44**<br>± **9.58** | **12.14**<br>± **7.83** | **4.630** | **0.001** |

TSP: Total Sleep Period; TST: Total Sleep Time; N1: NREM sleep stage 1; N2: NREM sleep stage 2; N3: NREM sleep stage 3; REM: Rapid Eye Movement sleep stage

However, individual metyrapone memory enhancement for the reactivated vs. non-reactivated story was not correlated with the above-mentioned individual sleep changes due to metyrapone (see Table S2).

## Discussion

This study shows that the reactivation of an episodic memory with a reminder cue at 3:55 a.m. followed by pharmacological cortisol suppression during early morning sleep enhances memory recall for the reactivated but not the non-reactivated story, tested four days after reactivation. Crucially, memory for the reactivated story was not only enhanced compared to the non-reactivated story in the cortisol suppression condition, but also in comparison to both the reactivated and the non-reactivated story in the placebo condition (reactivation by pharmacological manipulation interaction). Moreover, individual memory enhancement for the reactivated vs. non-reactivated story due to cortisol suppression was negatively correlated with the individual cortisol decrease during sleep after metyrapone, i.e. the more cortisol levels were suppressed during early morning sleep, the higher the increase in later memory for the reactivated vs. non-reactivated story.

The novelty of this study is a specific enhancement of a briefly reactivated episodic memory in humans likely as a consequence of altered reconsolidation processes due to cortisol suppression applied immediately after memory reactivation. Our study closely followed set criteria to test for pharmacologically-induced changes of reconsolidation (*8, 11*): (a) a consolidated memory was reactivated by a reminder cue, (b) the substance (manipulation) was administered after reactivation, and (c) memory was tested more than 24 hours later. Considering criterion (a), here, a 3-day-old and therefore consolidated memory was reactivated with a reminder cue (presentation of the first photo of the story and photo-related questions). As expected, at the time of memory reactivation (3:55 a.m.), prior to substance administration, baseline cortisol levels did not differ between the placebo vs. metyrapone condition. As set out in criterion (b), following memory reactivation, 3g of metyrapone were administered at 4 a.m., i.e. the pharmacological manipulation aimed at altering reconsolidation was applied post-reactivation. This pharmacological intervention suppressed cortisol levels, in particular, the morning cortisol peak, as depicted by significantly lower salivary cortisol levels in the metyrapone vs. placebo condition at all measurement points from 6:45 to 9:45 a.m., replicating previous findings (*35, 36*). As such, our pharmacological manipulation lowered cortisol levels for at least six hours after memory reactivation. To allow a sufficiently large time window for reconsolidation to take place, in accordance to criterion (c), memory for both the re-activated and the non-reactivated story were tested after four additional days (*34, 39*). Having observed these criteria, our finding that cortisol suppression specifically boosts memory for the reactivated story suggests that cortisol levels critically modulate memory reconsolidation processes of episodic memories.

This finding adds to our knowledge on episodic memory reconsolidation in humans. Previous studies using the same stimulus material showed memory to be impaired for the reactivated vs. non-reactivated story if propofol (medication that induces general anesthesia) or electroconvulsive shock therapy followed memory reactivation and memory was tested 24 hours later. Possibly, both manipulations led in a physiological blockade of episodic memory reconsolidation resulting in a later memory impairment (*11, 14*). In contrast, here we showed the opposite effect: Cortisol suppression boosted memory for the reactivated story, i.e. our pharmacological change in cortisol levels likely enhanced reconsolidation processes. Moreover, here, individual metyrapone-induced memory enhancement for the reactivated (vs. the non-reactivated) story, i.e. the source of the reactivation by manipulation effect, was negatively correlated to the individual cortisol decrease due to the pharmacological manipulation during sleep, indicating a direct relation of the two measures.

Interestingly, the main finding of this study contrasts previous literature on cortisol suppression effects on memory retrieval (*35, 40*), where participants showed impaired memory recall, when asked to recall their memories at a time when cortisol levels are acutely suppressed, i.e. metyrapone is already active (*35, 36, 40*). This recall impairment persists when tested a week later when cortisol levels are back to normal levels (*36*). These findings together with the current findings suggest that it is crucial whether a memory is retrieved under normal or under suppressed cortisol levels to influence later memory recall. If cortisol levels are low at the time of recall, acute and later memory recall are impaired, with metyrapone potentially altering acute memory recall as well as in subsequent reconsolidation processes. In contrast, if metyrapone is in-active at the time of reactivation and only administered thereafter as in the present study, metyrapone likely only affects reconsolidation processes, which may lead to altered outcomes such as enhanced memory, as found in the present study.

In addition, the effect of cortisol suppression altering reconsolidation is likely to depend on whether reconsolidation takes place during sleep or awake state. In a previous study in which we administered half the dose of metyrapone (1.5 gr instead of 3 gr as in the present study) at 9 a.m. (vs. 4 a.m. in the present study) after reactivation of one of the stories (as in the present study), we found memory to be decreased in the metyrapone vs. placebo condition independent of memory reactivation (*13*). This finding accords with studies on memory consolidation reporting that pharmacological administration of cortisol or cortisol synthesis inhibitor during sleep vs. wakefulness results in opposite effects on memory (*37, 38*). In addition, in our study metyrapone-induced cortisol suppression during early-morning sleep critically altered sleep architecture and quality/efficiency (increase in N1 and wake duration, and decrease in time spent in N3 and REM). However, we found no direct relations between individual changes in sleep due to metyrapone and memory enhancement due to metyrapone, an indication there might be a more complicated relation to be further investigated.

A limitation of our study is that we included only a small sample of women. Future studies should include a representative female sample to allow the generalization across genders of the reconsolidation effects of cortisol suppression. This is particularly important, as of now, female participants have not yet been tested in most of the studies examining metyrapone effects on memory (*35, 36, 40–42*). Additionally, future studies aiming to particularly address the role of sleep on cortisol-, stress-, or metyrapone-induced modulation of reconsolidation processes, should assess sleep parameters with extra caution and not as supplementary measures (as it was the case for this study), in order to be able to perform further analyses to a full sample of participants.

Altogether, this study shows that suppressing cortisol during early morning sleep alters standard reconsolidation processing and enhances memory for the material reactivated prior to the manipulation. This finding indicates that metyrapone-induced cortisol suppression actually reverses what may be the physiological function and effect of normal early-morning cortisol peak and respective sleep patterns to memory processing. Reactivation of past memories in early morning hours, physiologically followed by cortisol increase and REM sleep, seems to hinder their reconsolidation, in accordance to the described memory pruning function of sleep (*32, 33, 43, 44*). By contrast, reactivation of past memories in early morning hours, with pharmacological suppression of the morning cortisol peak as well as a consequent increase in time spent awake and decrease of N3 and REM sleep, rather enhances their reconsolidation, as shown in the present study. This finding may have strong implications for better understanding the phenomenon described as “over-consolidation” of memories in the case of post-traumatic stress disorder (PTSD; *33, 37, 45*), a disorder that has been associated with physiologically decreased cortisol levels.

## Materials and Methods

### Participants

Twenty healthy subjects (mean age 26 ± 4.67 years; mean body mass index 22.40 ± 2.4 kg/m^2^; 4 females) participated in this double-blind, within-subject cross-over study. They were free of neurological, psychiatric and endocrine disorders, not receiving any medication for the period of their participation (except for two of the women taking oral contraception), non-smokers, and free of any contraindication for metyrapone administration. All participants reported having a regular sleep-wake rhythm and spent one adaptation night in the sleep lab. The study was approved by the local ethics committee. All subjects provided written informed consent and were paid for their participation. Two male participants were excluded for being outliers (i.e. deviated more than two standard deviations from the group mean) in memory performance in the placebo condition.

### Stimuli

During the Encoding Session, participants were presented with two stories per condition (metyrapone/placebo). Each story comprised 11 slides (7 neutral, 4 emotionally arousing) accompanied by an auditory narrative. Each slide was presented for 20 sec. Participants were shown two previously used stories (*11, 13, 14, 46*), as well as two additional stories, parallel in structure and presentation from our laboratory.

### Experimental Design

After an adaptation night in the lab, each participant was tested in two conditions (metyrapone vs. placebo), with the order of condition counterbalanced across subjects. Conditions were separated by an interval of at least ten days. Each condition comprised an Encoding Session, a Reactivation Session and a Retrieval Session (**Figure 1A)**. In the Encoding Session, participants were presented two stories. In the Reactivation Session, two days after encoding, participants slept in the lab (lights off at 11:00 PM) and were awakened at 3:55 a.m., when one of the two stories was reactivated (see Reactivation below for more details). Directly following reactivation, at 4:00 a.m., the cortisol synthesis inhibitor metyrapone (3g, HRA Pharma) or placebo was orally administered with a light snack (yogurt). Then, participants slept until 6:45 a.m., when they were awakened. At the Retrieval Session, four days after reactivation (seven days after encoding), participants were asked to complete a multiple-choice recognition memory questionnaire for each of the two stories in each condition (see Multiple-choice recognition memory task below for more details).

#### Reactivation

At the Reactivation Session, one of the two encoded stories was reactivated using the procedure of previous studies (*11, 13, 14*). Participants were shown the first slide of one of the two stories, partially masked by three black-and-white checker-board patterns (**Figure 1A**). Participants were asked a question about the masked part of the scene. After providing their answer, they were presented with the same scene with a smaller mask (covering a smaller part of the scene), and finally no mask, i.e. the answer to each question was progressively revealed. The other story was not reactivated.

#### Multiple-choice recognition memory task

At the Retrieval Session, participants were tested for their memory of the reactivated as well as the non-reactivated story using a multiple-choice recognition memory test (*11, 13, 14*). Participants were asked 3-5 questions per slide (amounting to a total of 40 questions per story) presented in the order of the original slide shows. Answers to the first slide of all stories were excluded from analysis given that the first slide had been used for reactivation for one of the two stories in each condition. Memory performance scores represent the percentage of correct answers to all questions.

### Hormonal measures

Throughout the Reactivation Session, salivary cortisol samples were collected with Sarstedt salivette tubes (Sarstedt, Rommelsdorf Germany) at 3:55 a.m. (i.e. just before pill administration), at 6:45, 7:00, 7:15, 8:30, 8:45, 9:00, 9:15, 9:30 and 9:45 a.m. (**Figure 1B**). Saliva samples were stored at −25 C until sent for analysis. Cortisol levels were analyzed using luminescence immunoassay (IBL, Hamburg, Germany) and inter- and intra-assay coefficients of variations were below 5%.

### Polysomnographic recordings

Whole-night polysomnographic (PSG) recording was collected for both experimental nights. PSG included electroencephalography (EEG, 11 electrodes were placed according to the international 10-20 system), electrooculography (EOG), and electromyography (EMG) (*47*). The PSG signal was recorded with a V-Amp recorder (Brain Products, Gilching, Germany). All recordings were sampled at 512 Hz and stored for later offline analyses. EEG recordings were referenced to contralateral mastoids (A1, A2) for the offline analyses. Due to technical issues during one of the two experimental nights, analyses of sleep recordings were only performed on participants with complete datasets (n = 11).

### Statistical analyses

#### Behavior

Memory performance in the multiple-choice recognition memory task was analysed with a 2 (metyrapone/placebo) × 2 (reactivated/non-reactivated) mixed-design analyses of variance (ANOVA). Greenhouse-Geisser corrections of degrees of freedom were used when suitable and significant ANOVA effects were followed by pairwise t-test contrasts in order to specify the observed effects

Using correlation analyses, we further examined whether the individual suppression of the morning cortisol peak in the second half of the night (4:05 a.m. to 6:45 a.m.) and after waking up (6:45 a.m. to 9:45 a.m.) after metyrapone compared to the placebo condition, is related to the change in memory performance for the reactivated vs. non-reactivated story in the two conditions. In particular, a difference score for memory performance of the reactivated story minus memory performance for the non-reactivated story was calculated for each condition. The resulting difference score in memory performance between reactivated vs. non-reactivated story in the placebo condition was then subtracted from the corresponding difference score in the metyrapone condition score. This score will be referred as metyrapone memory enhancement for the reactivated vs. non-reactivated story, i.e. metyrapone memory enhancement = [(Reactivated – Nonreactivated Memory Performance)_Metyrapone condition_ – (Reactivated – Nonreactivated Memory Performance)_Placebo condition_]. To examine the changes of cortisol levels (morning cortisol peak in placebo vs. cortisol suppression after metyrapone condition) difference scores were calculated for cortisol level changes during sleep (cortisol level_6:45am_ - cortisol level_3:55am_) and after wakening (cortisol level_9:45am_ - cortisol level_6:45am_) for each condition. Then the corresponding cortisol change in the placebo condition was subtracted from the metyrapone condition (from now on referred as cortisol decrease due to metyrapone <u>during</u> sleep, i.e. [(cortisol_6:45am_ - cortisol_3:55am_)_Metyrapone condition_ - (cortisol_6:45am_ - cortisol_3:55am_)_Placebo condition_] and cortisol decrease due to metyrapone <u>after</u> sleep[(cortisol_9:45am_ - cortisol_6:45am_)_Metyrapone condition_ - (cortisol_9:45am_ - cortisol_6:45am_)_Placebo condition_]. We then correlated metyrapone memory enhancement with the change in cortisol decrease due to metyrapone during sleep (and after sleep respectively). We used Kendall’s tau b for these correlations, as more suitable to describe relations in smaller sample sizes (*48, 49*).

#### Cortisol levels

For the analysis of cortisol levels, separate linear mixed models were used (fitlme, MATLAB), in an effort to tackle missing values of cortisol levels (due to missing saliva samples, insufficient saliva quantity for the analyses, or cortisol levels below the assay’s sensitivity after metyrapone administration). Cortisol levels were log transformed to approach normal distribution of the residuals (note that untransformed cortisol levels are depicted at Figure 1B for illustration purposes). The linear mixed model for cortisol levels was set with fixed effects of factors substance (placebo/metyrapone) and time (10 time-points of the saliva samples/condition) and random effects of the factor subject. The marginal effects of factors substance and time were assessed with a type-III F-test, with the Satterthwaite approximation for the degrees of freedom, which is equivalent to omnibus repeated-measures ANOVA.

#### Sleep Analysis

Sleep analyses were conducted using PRANA software (Version 10.1. Strasbourg, France: Phitools). An expert scorer blind to the experimental conditions determined the different sleep stages (NREM 1, 2, 3, REM sleep and wake) for each recorded night of sleep. From the scoring of the sleep architecture, we computed the duration (min) of each sleep stage, as well as the percentage of each sleep stage relative to the total sleep period (TSP; from sleep onset to wake up time) and relative to the total sleep time (TST; TSP minus intra-wake epochs) for each phase of the night (i.e., sleep before the substance administration, sleep after the substance administration). Sleep efficiency (TST/time in bed*100) for each phase was also calculated. All extracted parameters were compared between metyrapone and placebo condition with pairwise t-test contrasts in order to identify differences in the sleep patterns between the two conditions. All the t-tests reported were two-tailed and for all analyses the significance level was set to *p* ≤ .05.

## Supplementary Materials

Results

Table S1. Output of linear mixed model on cortisol levels.

Table S2. Non-parametric correlations between memory and sleep parameters.

## General

This study was conducted at the Brain and Behavior Lab (BBL) and benefitted from support of the BBL technical staff.

We thank Virginie Sterpenich for training experimenters on the set-up of polysomnographic recordings. We thank Megan Mulligan, Florian Ruiz, Laura Riontino, Emalie McMahon, Anais Chevalley, Shannon Wilfred, and Vassilis Kehayas for help with data acquisition and analysis.

## Funding

This research was supported by grants from the Pierre Mercier foundation (to U.R.), the Swiss National Science Foundation (PZ00P1_137126 and PZ00P1_160861 and PCEFP1_186911), the European Community Seventh Framework Program [FP7/2007-2013] under grant agreement 334360, and the National Center of Competence in Research (NCCR) Affective Sciences and the Academic Society of Geneva (Foremane Fund). D.A. was supported by the Ernst and Lucie Schmidheiny Foundation.

## Author contributions

D.A. and U.R. designed the study. D.A. acquired the data. D.A. and U.R. analyzed memory and cortisol data; A.A.P. analyzed sleep data. D.A., U.R., A.A.P., S.S., and P.V. wrote the manuscript.

## Competing interests

There are no competing interests.

## Supplementary Materials

**Table S1.**
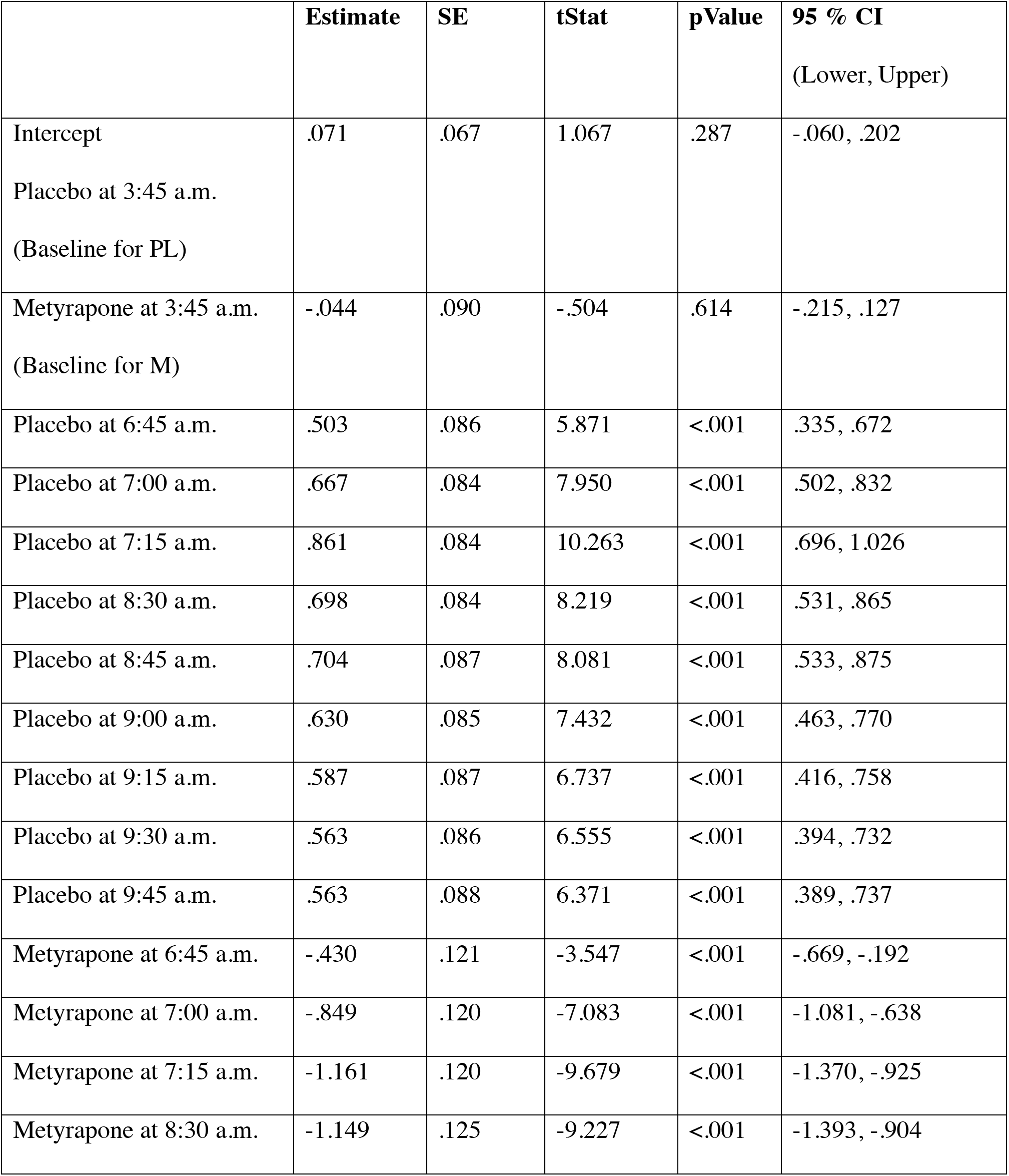
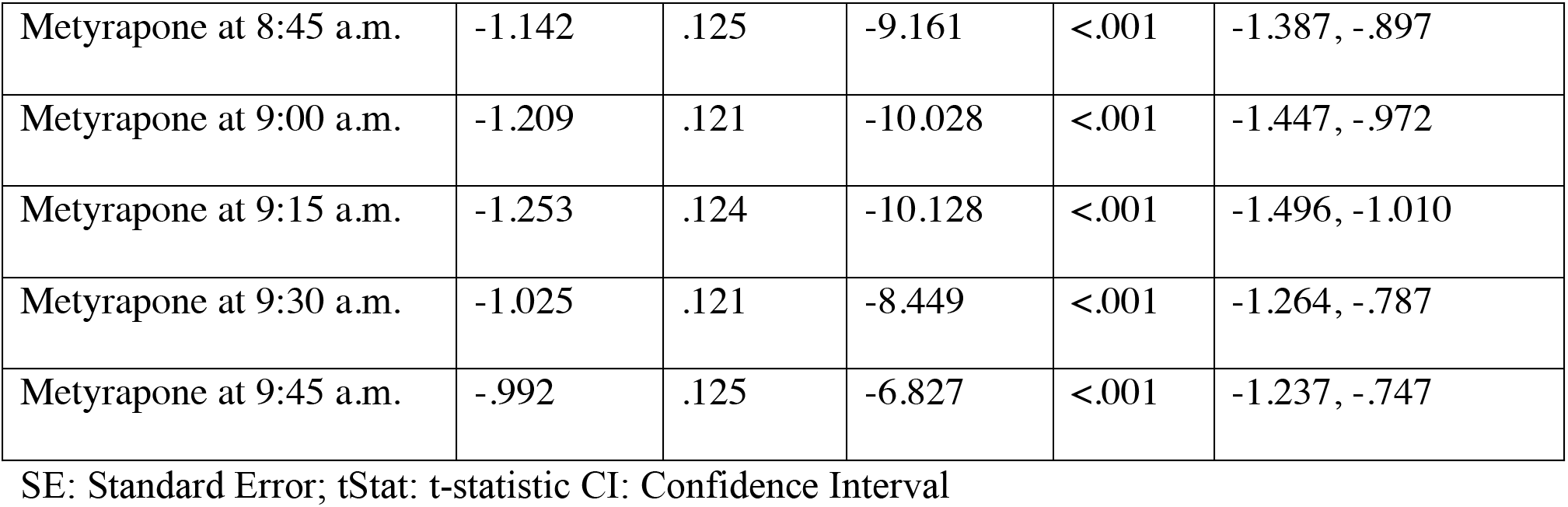
Output of linear mixed model on cortisol levels with fixed effects of factors treatment (placebo/metyrapone) and time (10 time-points of the saliva samples/condition) and random effects of the factor subject.

**Table S2.**
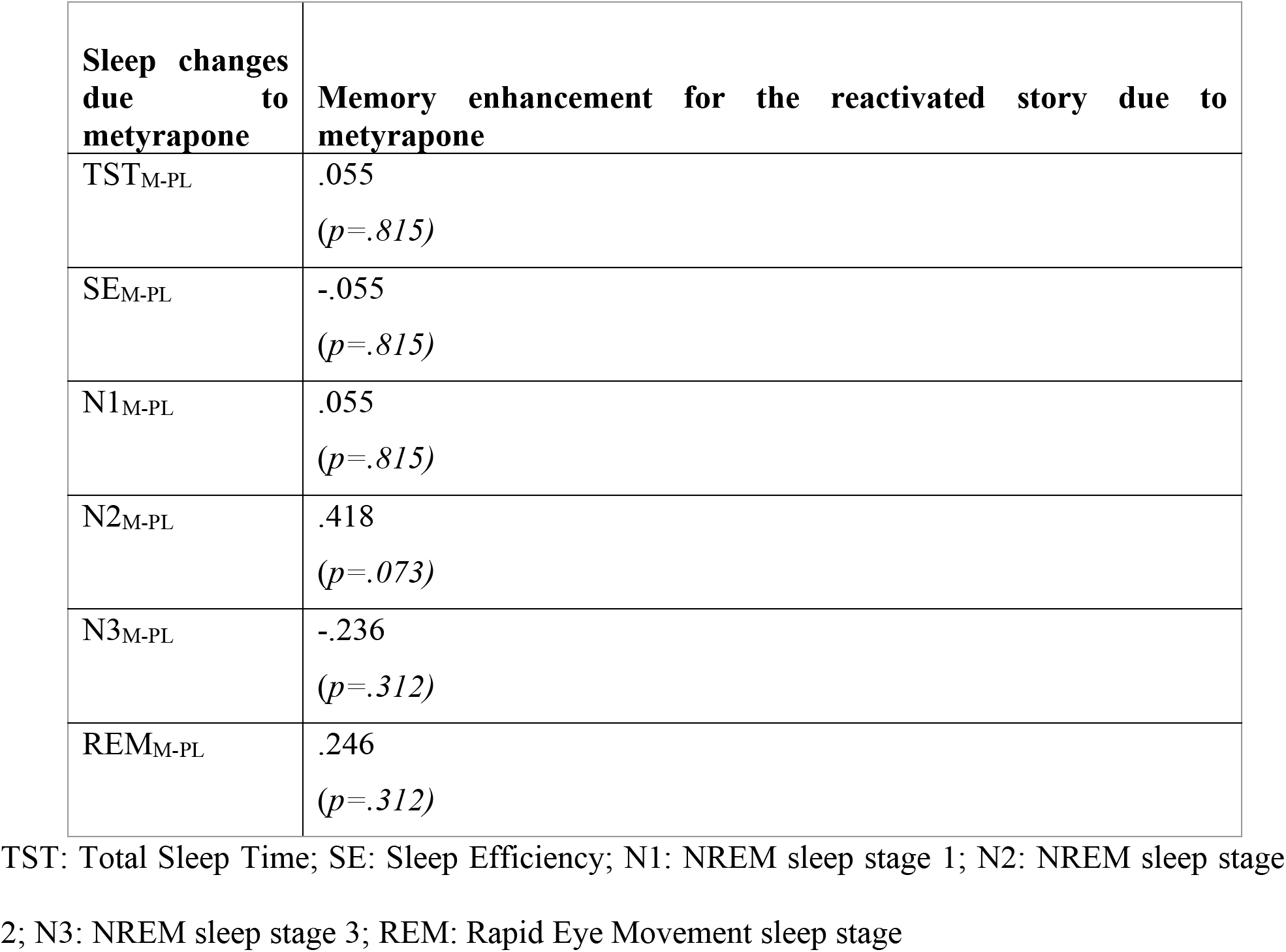
Non-parametric correlations between individual me metyrapone memory enhancement for the reactivated vs. non-reactivated story and changes in sleep parameters due to metyrapone.

## Notes

### Competing Interest Statement

The authors have declared no competing interest.

